# *LINC00261* expression reduces mutation accumulation while conferring resistance to cisplatin and altering the tumor-immune microenvironment in lung adenocarcinoma

**DOI:** 10.1101/2023.03.02.530823

**Authors:** Jonathan Castillo, Tianchun Xue, Samantha Joseph, Mayela Norwood, Daniel J. Mullen, Stephanie Berrellez, Sophia James, Alan L. Epstein, Troy McEachron, Crystal N. Marconett

**Author notes:** Correspondence and requests for reprints should be addressed to Crystal Marconett, Ph.D., Norris Comprehensive Cancer Center, Keck School of Medicine, University of Southern California, NRT 2504, Los Angeles CA 90033-9520, USA.

## Abstract

Immunotherapy has emerged as a major breakthrough that has the ability to improve survival outcomes for multiple solid tumor types, however effective response requires the combination of blocking inhibitory signals on the tumor surface and antigen presentation to the tumor surface for proper immune recognition. Several commercially available and robust methods exist for identification of tumors displaying immune-inhibitory surface receptors, including PD-L1 and CTLA-4, however it is currently difficult to predict effectiveness of antigen presentation on the cell surface. To address this, we utilized clinical and next-generation sequencing data generated by The Cancer Genome Atlas (TCGA) to identify gene signatures that are correlated to tumor mutational burden (TMB) within cancers of epithelial origins as a surrogate for neoantigen signatures. We identified *LINC00261* as a top gene correlated to mutational burden (TMB), whose expression activates DNA damage response pathways *in vitro* along with resistance to cisplatin. *LINC00261* expression was also significantly correlated to MHC Class II genes involved in endogenous neoantigen presentation expression within the TCGA-LUAD cohort. This relationship was confirmed *in vitro* through ectopic reintroduction of *LINC00261* for key MHC Class II presentation genes. Interferon gamma-induced MHC gene activation *in vitro* was also able to induce endogenous expression of *LINC00261*. Our results suggest there is a mechanistic relationship between *LINC00261* expression and functional MHC Class II neoantigen presentation in LUAD. The silencing of *LINC00261* in LUAD results in compromised DNA repair, accumulation of mutations, and reduced antitumor immune response.

## 1. Introduction

One of the major driving forces of carcinogenesis is the accumulation of somatic DNA mutations. Rates of mutation vary considerably among tumor types, with malignant melanoma and lung adenocarcinoma among the highest rates of DNA mutation accumulation [1–3]. In lung cancer, this is due in part to exposure from environmental mutagens, such as tobacco smoking and air particulate pollution. Accumulation of mutations can also be attributed to loss of effective DNA damage response (DDR) [4,5].

Total mutational burden (TMB) is used clinically in conjunction with PD-L1 and CTLA4 status to determine which patients may benefit from immunotherapy [6]. TMB is a relevant clinical marker due to being a surrogate for neoantigen load, which is critical for anti-tumor immune recognition and response [7,8]. Understanding the key mechanisms causing elevated TMB can give us a molecular understanding of the basis for the efficacy of immune checkpoint therapies. This strategy has to date been highly successful in melanoma, lung cancers, and other solid tumors, with moderate success seen in solid tumors of endodermal origin [6]. Specifically, in non-small cell lung cancer (NSCLC), TMB predicted better response to PD-1/CTLA-4 inhibitors, with patients who experienced complete/partial response having higher TMB compared to those with stable or progressive disease (median 273 versus 114 mutations, respectively, Mann-Whitney p-value of 4.0×10^-4^) [9].

Understanding the molecular basis of which patients may experience therapeutic benefit is paramount to improving clinical outcomes. Although deficiencies in expression and function of some DDR genes correlate with high TMB levels, recent studies have suggested that DDR gene alterations themselves can serve as predictive biomarkers of immunotherapy response [10]. In addition to immunotherapies, platinum-based chemotherapeutics have also shown efficacy towards patients with tumors containing high TMB [11]. Determining which patients are most likely to respond prior to chemotherapeutic administration could significantly improve patient outcomes.

Although progress has been made in the development of patient response to immune checkpoint inhibitors (ICI), there are patients who still fail to respond, even with predicted response markers such as high TMB [12,13]. With the expansion of transcriptional profiling, non-coding RNAs have emerged as major regulators of carcinogenesis. Long non-coding RNA (lncRNA) are transcripts greater than 200 nucleotides in length with little to no coding capability that are increasingly recognized as key regulators of signaling pathways [14]. Dysregulation of lncRNA *HOXA11-AS* [15] and *ANRIL* [16], among other lncRNAs, have been recently characterized as drivers of cisplatin-resistance [17]. LncRNAs significantly associated with TMB but without characterized functions could provide an opportunity to understand the molecular underpinnings of patient response to chemotherapeutics, however, to date no systematic study has determined the extent of lncRNA-TMB relationship in cancer.

Here we evaluate the connection between lncRNA function, DNA damage repair, and how these affect patient response to chemotherapy in lung adenocarcinoma. Previous work has identified *LINC00261* as a tumor suppressor in lung adenocarcinoma that acts as a critical mediator of the DDR [18] however the relationship between *LINC00261* and therapeutic response was unknown. Our results indicate that several lncRNAs are associated with TMB, with *LINC00261* being one of the top correlated lncRNAs in LUAD. In-depth mechanistic characterization of *LINC00261* demonstrated a clear functional link between this lncRNA and TMB as well as subsequent neoantigen presentation and was able to affect LUAD cellular response to platinum-based chemotherapeutics.

## 2. Materials and Methods

### 2.1. Cell Lines, Culture Conditions, and treatment conditions

The human LUAD cell lines HCI-H522 and A549 were obtained from the laboratory of E. Haura and the American Type Culture Collection (ATCC), respectively. Cell lines were fingerprinted at the University of Arizona Genetics Core to verify their identity prior to experimentation. The monoclonal stably transfected cell lines A549-shScrambled, A549-*shLINC00261* H522-CMV-NEO, and *H522-CMV-LINC00261* were previously generated in triplicate, as described in Shahabi et al., [18]. Cell lines were cultured in RPMI-1640 medium (Corning, Cat# 45000-396) containing 10% fetal bovine serum (X&Y Cell Culture, Cat# FBS-500, Lot# 7B0302) and 100 U/mL penicillin/streptomycin (VWR Life Sciences, Cat# 82026-730). For IFNγ (R&D Systems, Cat# 285-IF-100/CF) treatment, cells were treated with 100 ng/mL for 24 hours.

### 2.2. Cytotoxic assay

Cell viability was determined using a Cell Counting Kit-8 assay (Abcam, Cat# ab228554). Cisplatin (Enzo Life Sciences, Cat # ALX-400-040-M050), carboplatin (Sigma, Cat# C2538), and oxaliplatin (Sigma, Cat# O9512) were dissolved in dPBS (Corning, Cat# 45000-434). Paclitaxel (Tocris Bioscience, Cat# 109710) was dissolved in DMSO (Sigma, Cat# D8418). A day prior to chemotherapeutic treatment, A549 and H522 cells were seeded on 96-well plates at 3000 and 5000 cells per well, respectively. Three days after incubation, cells tested for viability using Cell Counting Reagent 8 (Abcam, Cat# ab228554). Briefly, 1:9 by volume of CCT-8 to complete-culture medium was added per well and incubated for 4 hours. Viability was measured using Multiskan FC microplate photometer (ThermoFisher Scientific, Cat# 51119000) at absorbance of 460 nm. Percent cell viability was normalized to no treatment controls. Cytotoxic assays were performed with three biological replicates, i.e. three different monoclonal stable transfected cell lines.

### 2.3. RNA isolation and qRT-PCR

Total RNA was isolated and purified from cultured cell lines using the Aurum Total RNA Mini kit (Bio-Rad, Cat# 7326820). RNA was converted to cDNA using iScript cDNA Synthesis Kit (Bio-Rad, Cat# 1708891) and qPCR was performed using SYBR Green Supermix (Bio-Rad, Cat# 1708886), following manufacturer’s protocol. Reaction steps consisted of an initiation step at 95° C for 3 minutes, followed by 50 cycles of denaturation for 30 seconds at 95° C, annealing for 30 seconds at 57° C, elongation for 30 seconds at 72° C, and plate reading. Relative expression was calculated using the double delta Ct method, where GAPDH was used as the internal control gene. Three technical replicates were done for each qPCR reaction and Ct were averaged. Number of biological replicates is indicated in the figure legend. PCR primer sequences are described in Supplementary Table 1.

### 2.4. High dimensional data analysis

TCGA (The Cancer Genome Atlas) data (Data Release 34.0 - July 27, 2022) of endocervical adenocarcinoma (CESC), esophageal carcinoma (ESCA), stomach adenocarcinoma (STAD), lung adenocarcinoma (LUAD), bladder urothelial carcinoma (BLCA), uterine corpus endometrial carcinoma (UCEC), skin cutaneous melanoma (SKCM), head and neck squamous cell carcinoma (HNSC), lung squamous cell carcinoma (LUSC), and colorectal adenocarcinoma (COAD) were downloaded from the GDC (Genomic Data Commons) repository using the “TCGAbiolinks” package (version 2.16.0) [19] in R (version 4.0.0). TMB was obtained from the TCGA mutation format (MAF) files. The “stringr” package was used to substring the barcodes of MAF and expression data (FPKM-UQ). Log2 fold change of expression data of tumor samples was used to calculate the Pearson correlation to tumor TMB levels, and false discovery rate (FDR)-corrected p-values were applied to account for multiple comparisons being performed. Significantly correlated genes were selected by setting an FDR-corrected p-value cutoff ≤ 0.05. The positive or negative value of Pearson correlation coefficient was used to determine the positive or negative relationship of the correlation. LncRNA versus mRNA classification was annotated using GENCODE (gencode.v34.LncRNAs) and imported in R by the “rtracklayer” package (version 1.47.0) [20]. Heatmaps were generated with the “Pheatmap” package (version 1.0.12) [21] in R, using ward.D2 as the clustering method and colored by their correlation values and gene types. Specifically, genes included in the heatmap were significantly correlated to TMB in LUAD and at least one other cancer type among the ten cancer types tested.

### 2.5. Common Gene Pattern Analysis

Genes expressed in both LUAD and STAD were selected and their Pearson correlation to TMB was calculated to determine if the two cancers have similar TMB-related gene patterns. Log10-transformed FDR-corrected p-values were used to indicate the significance of correlation, with a significance cutoff of ±1.3. Adjusted p-values of genes in LUAD and STAD were shown as a starburst plot generated by the “ggplot2” package (version 3.3.0) [22] in R, colored by their different types of correlation.

### 2.6. Correlation Analysis

Smoking status was obtained from TCGA-LUAD clinical data. Smoking status “1” (non-smokers) was defined as never smokers, “2” (current smokers) was classified into current smokers, “3” (Current Reformed Smoker for > 15 years) & “4” (Current Reformed Smoker for <=15 years) were classified as former smokers. Samples were divided into *LINC00261*-expressing or silenced groups at FPKM of 1. Box plots were generated with ‘ggplot2’ (version 3.3.0) in R [22].

### 2.7. Pathway Analysis

QIAGEN Ingenuity Pathways Analysis (version 01-12) was used to determine pathway enrichment with Benjamini-Hochberg correction on significantly correlated mRNAs (BH-corrected p-value ≤ 0.05) for the statistical likelihood of enrichment. TMB-correlated genes underwent pathway enrichment analysis, using Qiagen Ingenuity Pathways Analysis (IPA) [23]. Bar plots were generated by the “ggplot2” package (version 3.3.0) [22] in R as well as to visualize the enriched pathways with a significance threshold of −log 10 BH corrected p-value set to 1.3 (-log10 BH of 0.05).

### 2.8. Analysis of mutational signatures and mutational patterns

Identification of significantly present mutational signatures was done by using the R package The Mutational Signature Comprehensive Analysis Toolkit (musicatk) [24]. Then the extracted signatures were compared with COSMIC signatures and visualized. To classify single nucleotide variants (SNVs) in LUAD MAF files into transition and transversions mutations, the maftools package (version 2.4.12) [25] in R was used. SNV was visualized as a box plot showing the overall distribution of six different mutation types. Then, TCGA-LUAD samples were subset into two groups based on *LINC00261*-expressing (FPKM > 1) or silenced (FPKM ≤ 1), and the fraction of individual mutation types was then compared between the two groups. Box plots were generated by “ggplot2” (version 3.3.0) in R [22]; and an unequal variances (Welch) t-test was used to determine significance (p-value ≤ 0.05).

## 3. Results

### 3.1. Genome-wide correlation of gene expression and mutational burden within epithelial cancers reveals association of LINC00261 to TMB

To determine the impact of lncRNA association with the accumulation of DNA mutations in cancer, we utilized whole exome sequencing and patient-matched transcriptomic profiling available from The Cancer Genome Atlas (TCGA) [26]. We identified genes associated with mutational accumulation by performing a transcriptome-wide Pearson correlation between total tumor mutational burden (TMB) and the expression level of individual genes, which included both protein-coding mRNAs and polyadenylated lncRNAs (Figure 1a). Among correlated genes within the 10 cancer types, 62.95% were negatively correlated to TMB and 37.04% were positively correlated to TMB (Figure 1b,c). We observed that 598 mRNAs and 93 lncRNAs were associated with TMB in both LUAD and another solid cancer type. Stomach adenocarcinoma (STAD) displayed the most consistent TMB-lncRNA relationship to LUAD (Figure 1a).

**Figure 1.**
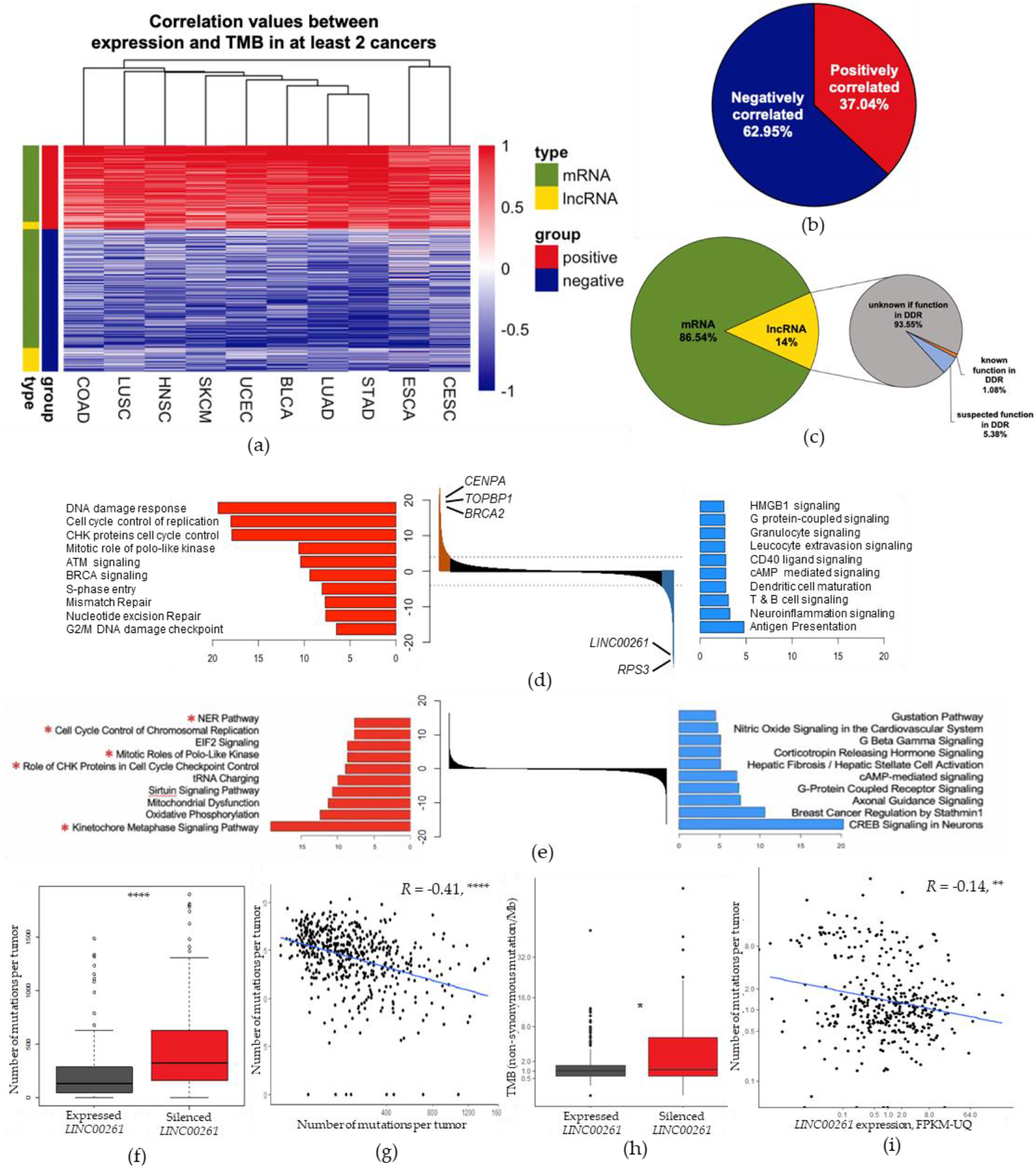
Genome-wide correlation of gene expression and mutational burden within epithelial cancers reveals association of *LINC00261* to TMB (a) Supervised clustering heatmap of correlation values between gene expression and TMB across cancer types. The depth of colors represented the level of correlation; red = positively correlated, blue = negatively correlated. Green = mRNA, yellow = lncRNA. (b) Pie chart indicating the percentage of genes that were positively (red) or negatively correlated (blue) to TMB. (c) Distribution of mRNA vs lncRNA genes correlated to TMB. Among lncRNA genes, information attributed to DDR function according to IPA functional annotation is noted. (d) TCGA-LUAD genome-wide correlations between gene expression and mutational burden. Genes are ranked by correlation of their transcriptional expression to mutational burden. Red = positive correlation, blue = negative correlation. Bar plot shows pathways enriched within the positively (red) or negatively (blue) correlated genes to TMB. (e) TCGA-STAD genome-wide correlations between gene expression and mutational burden. Genes are ranked by correlation of their transcriptional expression to mutational burden. Red = positive correlation, blue = negative correlation. Bar plot shows pathways enriched within the positively (red) or negatively (blue) correlated genes to TMB. (f) Boxplot of TMB in TCGA-LUAD patients based on *LINC00261* expression status. Silenced *LINC00261* is FPKM < 1). (g) Scatter plot of *LINC00261* expression (in FPKM-UQ) compared to TMB for each patient with both WES and expression data from TCGA-LUAD, n=502. (h) Boxplot of TMB in TCGA-STAD patients based on *LINC00261* expression status. Silenced *LINC00261* (cutoff FPKM < 1). (i) Scatter plot of *LINC00261* expression (in FPKM-UQ) compared to TMB for each patient with both WES and expression data from TCGA-STAD, n=368.

To ascertain mechanistic information on the genes that are correlated to TMB, we performed a pathway enrichment analysis using Ingenuity Pathway Analysis (IPA). After separating TMB-associated genes as either positively correlated (red) or negatively correlated (blue) (Figure 1d,e), we observed that there was a highly statistically significant correlation between DNA damage response (DDR) genes and TMB in LUAD. TMB-associated mRNA whose expressions were positively correlated to TMB were ascribed function in DDR, with pathways including DNA damage response, cell cycle control of replication, ATM signaling, being significantly enriched (a −log10 BH values greater than 1.5). This included TOPBP1, CENPA, and BRCA2, among other known genes involved in DDR [27,28]. Genes that were anti-correlated with TMB were associated with the immune response, such as genes involved in “antigen presentation”, “T and B cell signaling”, and “neuroinflammation signaling” and other pathways, all having a significant enrichment above −log10 BH p-value > 1.5 (Figure 1d). MHC class II expression, which may represent enrichment for antigen-presentation in cells such as dendritic and alveolar macrophages in tumors with elevated TMB.

The lncRNA *UBXN10-AS1* was among the top lncRNA correlated to TMB (FDR p-value 1.07e-14), followed by *LINC01550* (1.71e-11) (Table 1). Both lncRNAs have previously described roles as tumor suppressors in various cancers, with *UBXN10-AS1* inducing cell cycle arrest [29], and *LINC01550* associated with favorable immune microenvironment [30]. In contrast, *LINC00261* was the third ranked lncRNA correlated to TMB in LUAD, and strikingly has a previously described role in DNA damage response [18]. In the TCGA data cohort, *LINC00261* was significantly inversely correlated to TMB in LUAD (Spearman R: −0.41, p < 2.2 x 10^-16^) (Figure 1f,g). A similar observation was seen in STAD, in that lower expression of *LINC00261* correlates to higher TMB, compared to silenced *LINC00261* tumors, with a significant p-value of 0.013 (Figure 1h,i). This significant inverse relationship was also observed for *TP53TG1*, a previously identified lncRNA that alters p53 response to DNA damage (Supplementary Figure 1) [31,32].

**Table 1.** LncRNAs with the highest statistically significant negative correlation to TMB, in LUAD

| ENSG ID | Gene Name | p-value | FDR Corrected p-value | Pearson Correlation Coefficient |
| --- | --- | --- | --- | --- |
| ENSG00000225986 | *UBXN10-AS1* | 1.30E-17 | 1.07E-14 | -0.37011 |
| ENSG00000246223 | *LINC01550* | 8.42E-14 | 1.71E-11 | -0.32609 |
| **ENSG00000259974** | ***LINC00261*** | **8.96E-14** | **1.79E-11** | **-0.32575** |
| ENSG00000177133 | *LINC00982* | 1.50E-13 | 2.76E-11 | -0.32293 |
| ENSG00000281406 | *BLACAT1* | 4.14E-12 | 5.29E-10 | -0.30403 |
| ENSG00000266256 | *LINC00683* | 1.02E-11 | 1.14E-09 | -0.29863 |
| ENSG00000237248 | *LINC00987* | 1.14E-11 | 1.26E-09 | -0.29795 |
| ENSG00000245937 | *LINC01184* | 1.16E-11 | 1.27E-09 | -0.29789 |
| ENSG00000258498 | *DIO3OS* | 1.24E-11 | 1.35E-09 | -0.29746 |
| ENSG00000223392 | *CLDN10-AS1* | 1.90E-11 | 1.96E-09 | -0.29486 |
| ENSG00000225194 | *LINC00092* | 3.07E-11 | 2.96E-09 | -0.29191 |
| ENSG00000234456 | *MAGI2-AS3* | 3.48E-11 | 3.31E-09 | -0.29114 |
| ENSG00000268388 | *FENDRR* | 4.80E-11 | 4.33E-09 | -0.28913 |
| ENSG00000255399 | *TBX5-AS1* | 5.71E-11 | 4.97E-09 | -0.28804 |
| ENSG00000263812 | *LINC00908* | 6.37E-11 | 5.49E-09 | -0.28735 |
| ENSG00000230590 | *FTX* | 1.31E-10 | 1.01E-08 | -0.28277 |
| ENSG00000235049 | *LINC00940* | 1.68E-10 | 1.25E-08 | -0.28116 |
| ENSG00000225329 | *LHFPL3-AS2* | 2.53E-10 | 1.77E-08 | -0.2785 |
| ENSG00000267280 | *TBX2-AS1* | 2.75E-10 | 1.90E-08 | -0.27796 |
| ENSG00000225434 | *LINC01504* | 2.91E-10 | 1.99E-08 | -0.27759 |

### 3.2 Association of LINC00261 with clinical covariates and known DNA repair pathways

In LUAD, several clinical variables are associated with an increased susceptibility to mutation, such as smoking status and age at diagnosis [33,34]. To understand how these clinical variables correspond with *LINC00261* expression, a multiple linear regression model was fit to predict expression of *LINC00261* using clinical variable information available in TCGA, including sex, race, age of diagnosis, smoking status, and TMB (Table 2). *LINC00261* was significantly correlated to age of diagnosis in adjacent normal tissue, demonstrating a decrease in expression of *LINC00261* overtime (Figure 2a).

**Table 2.** Multiple linear regression analysis of LINC00261 expression in LUAD

|  | Estimate | p-value |
| --- | --- | --- |
| Intercept | 10.9397 | 0.000129*** |
| Gender |  |  |
| *Female* | -0.56221 | 0.451317 |
| Race |  |  |
| *Black* | -1.04549 | 0.361448 |
| *Asian/Pacific Islander* | 0.59063 | 0.832997 |
| *American Indian/Alaska Native* | -1.94286 | 0.790228 |
| Age at Diagnosis |  |  |
| *Year* | -0.09243 | 0.017684* |
| Smoking History |  |  |
| *Former Smoker > 15* | 1.91761 | 0.124998 |
| *Former Smoker ≤ 15* | -0.54069 | 0.657167 |
| *Current Smoker* | -0.7724 | 0.560219 |
| Mutation Status |  |  |
| *Tumor mutational burden* | -0.12521 | 0.02496* |

Multiple $R^2 = 0.04781$ , Adjusted $R^2 = 0.02623$ , p-value = 0.0204

**Figure 2.**
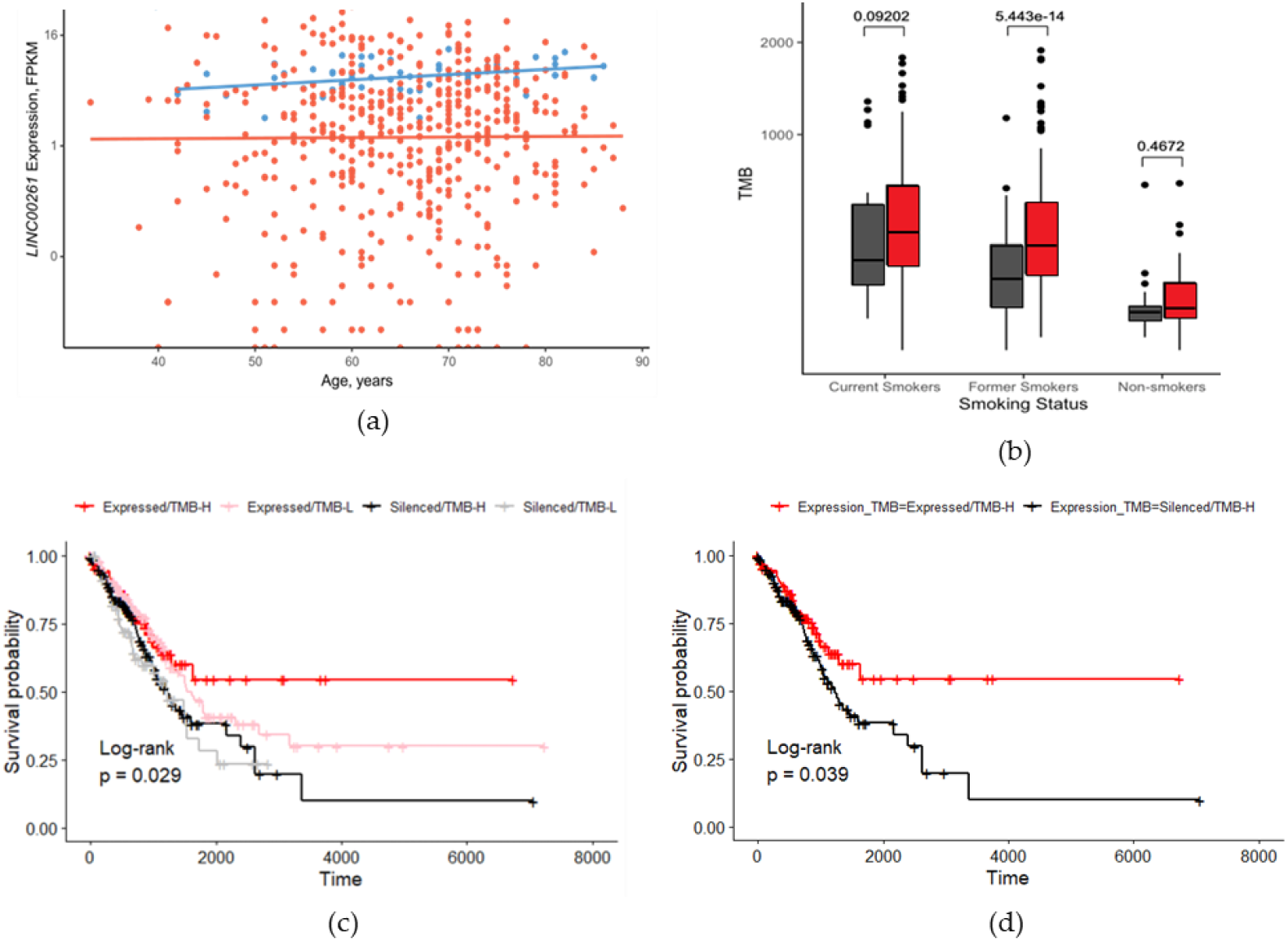
Loss of *LINC00261* expression is associated with several clinical variables in LUAD. (a) *LINC00261* expression of TCGA samples based on age. Blue - Adj. normal tissue, orange - primary tumor tissue. (b) Boxplot of LINC00261 expression compared to mutational burden, broken-down by smoking status. (c) TCGA-LUAD patients were stratified by expression status of *LINC00261* (expressed: FPKM-UQ > 1, silenced: FPKM-UQ ≤ 1) and TMB (TMB-H: above median, TMB-L: below median). Significance difference in survival outcome between groups was tested by log-rank test. (d) Survival outcome of patients with high TMB (TMB-H) between those with expressing-*LINC00261* and silenced-*LINC00261*.

In LUAD, the top contributor to mutational load is smoking [35]. As *LINC00261* was observed to be associated with the number of mutations in LUAD patients, we investigated whether *LINC00261* also influenced the mutational load due to smoking. To do this, the TCGA LUAD cohort was separated by smoking status into distinct categories: (1) never-smokers, (2) former smokers who quit more than 15 years ago, (3) former smokers who quit less than 15 years ago, and (4) current smokers. In the non-smoking cohort (“1”), we saw no significant difference in the number of mutations based on *LINC00261* expression status, with a p-value of 0.467 (Figure 2b). This was not the case in current smokers (“4”) or former smokers (“2”, “3”), as there was a trend of difference and a significant difference in the number of mutations in patients with silenced *LINC00261* with a p-value of 0.092 and 5.4×10^-14^, respectively (Figure 3b).

**Figure 3.**
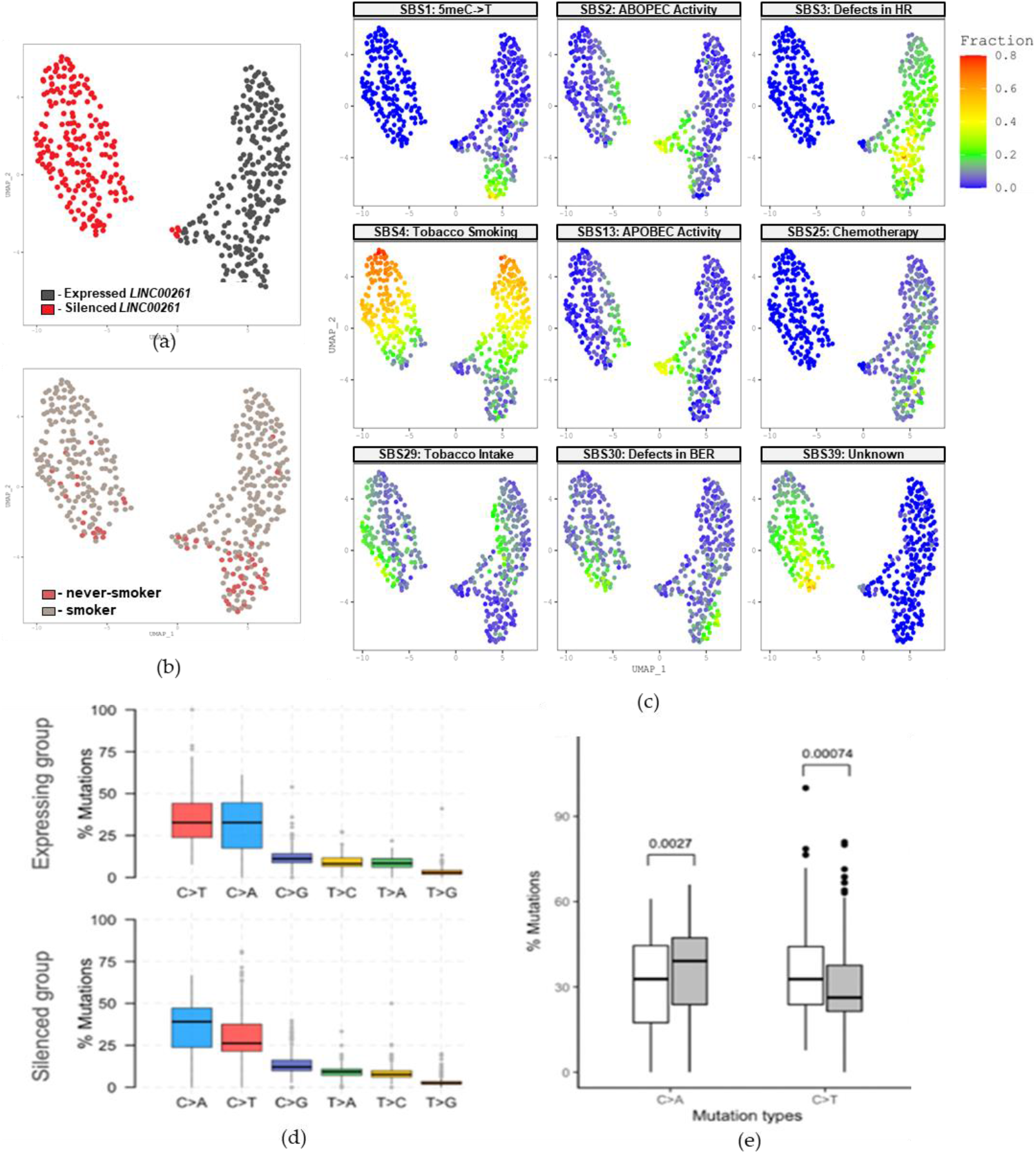
Mutational patterns present in patients with silenced *LINC00261* expression. (a) Clustering of TCGA-LUAD samples based on fraction of mutational signatures which were predicted to be significantly enriched within silenced-*LINC00261* and expressing-*LINC00261* sample sets. (b) Same as B but smoking status is labeled. (c) Presence of mutational signature found in each TCGA-LUAD patient sample. (d) Box plot showing the percentage of 6 types of mutation, including C>T, C>A, C>G, T>C, T>G, T>A. (e) Box plot showing the percentage of C>T and C>A mutations in *LINC00261-* expressing and silenced groups. White boxes are the *LINC00261*-expressing group, and gray boxes are the *LINC00261*-silenced group. An unpaired t-test was used to test for significant difference between the silenced and expressing g

### 3.3 Patients with silenced LINC00261 expression demonstrate different mutational patterns

In order to delineate the mechanism by which silencing of *LINC00261* resulted in higher mutational burden, we sought to identify the mutational signatures present in patients with silenced *LINC00261* expression. Mutational signatures are patterns of somatic mutations arising from carcinogenic exposures or dysregulated repair mechanisms [36]. By identifying the mutational patterns present within our cohort of study, said patterns can be compared to signatures generated experimentally to infer on the source of the mutagenic events due to silenced *LINC00261* [5,37,38]. We clustered TCGA-LUAD samples based on fraction of mutational signatures which were predicted to be significantly enriched within silenced-*LINC00261* and expressing-*LINC00261* samples sets (Figure 3a). Because smoking is a major source of mutations, smokers and never smokers were highlighted (Figure 3b). Patients with silenced *LINC00261* expression demonstrated a higher number of mutational features associated with tobacco uptake (SBS4, SBS29), defects in BER (SBS30), and other repair mechanisms (SBS39) (Figure 3c, Supplemental Figure 2), demonstrating that silenced *LINC00261* correlates with increase damage from these mutagenic events. In addition to the difference in mutational signatures present in TCGA-LUAD patients with silenced *LINC00261* we also noticed differences in the overall base pair substitution types (Figure 3d). Patients with silenced *LINC00261* had higher incidence of C>A transversion (p = 0.0027) and lower C>T transitions (0.00074), respectively (Figure 3e). The C>A transversion has previously been observed in smokers [39].

### 3.4. LINC00261 loss is associated with inactive DNA repair and sensitivity to cisplatin

In order to determine the relationship between *LINC00261* and DNA repair mechanisms, we looked at whether *LINC00261* expression correlated with dysfunctional DDR pathways. This was done by identifying if a gene within a given DDR pathway underwent a loss of function alteration, such as a functionally deleterious mutation, copy number deletion, or DNA methylation silencing. Genes associated with a given DDR pathway were sourced from previously curated lists and are also available in Supplementary Table 2 [40,41]. Previously curated values for whether a DDR gene within a given TCGA patient had a loss of function alteration was collected from TCGA DNA Damage Repair Analysis Working Group (DDR-AWG) [40]. Briefly, the DDR-AWG is among the ten AWGs involved in the TCGA PanCancer Atlas project, where the DDR-AWG used newly standardized Pan-Cancer Atlas data to systematically analyze potential causes of loss of DDR function [40,42]. We observed that *LINC00261* expression is significantly lower in patients with loss of function alterations in nucleotide excision repair (p-value of 1.8×10^-5^), base excision repair (p-value of 3.3×10^-7^), and other repair pathways, as seen on Figure 4a.

**Figure 4.**
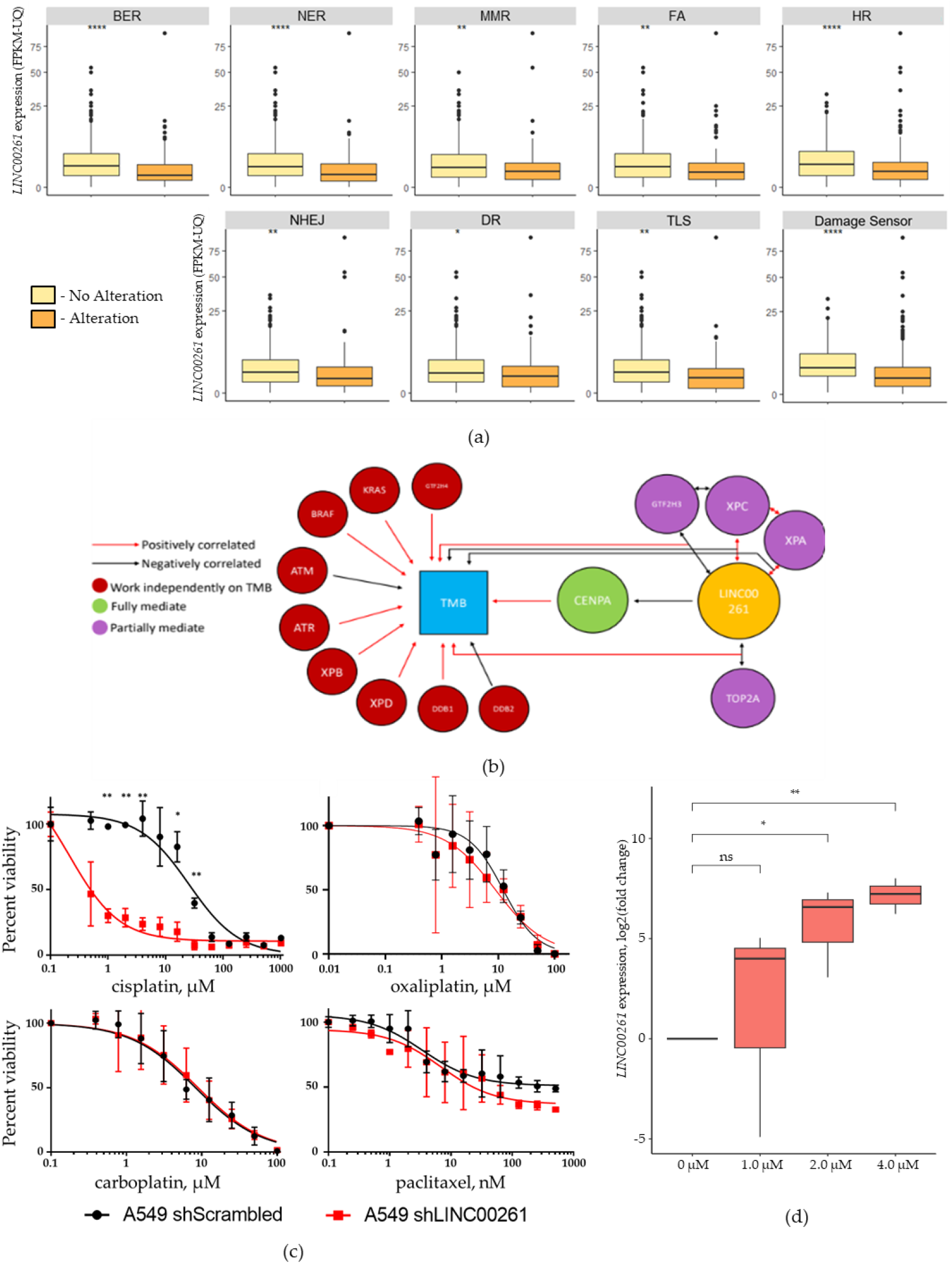
*LINC00261* expression association with DNA damage response pathways and susceptible to DNA damaging agents. (a) Expression of *LINC00261* in TCGA-LUAD samples with (orange) or without alterations (yellow) in the respective DDR pathway. Patients were classified as having an altered pathway if a gene within a given DDR pathway underwent a loss of function alteration, such as a functionally deleterious mutation, copy number deletion, or DNA methylation silencing. (b) Diagram of mediation analysis between factors and TMB. The arrows represent two factors correlated to each other. Red arrow represents positively correlated; black represents negatively correlated. (c) Cell viability of A549-sh*LINC00261* (red) compared to A549-shScrambled (black) controls when exposed to the DNA damaging agent cisplatin, paclitaxel, carboplatin, or oxaliplatin. (d) RT-qPCR of changes in *LINC00261* expression when treated with cisplatin for three days, compared to no treatment vehicle control (saline). An unpaired t-test was used to test for significant difference between no treatment (0 μM) and treatment conditions, with three replicates.

To understand the causal sequence by which loss of DDR function, *LINC00261* downregulation, and TMB arise in LUAD, we performed a mediation analysis. A linear model was used to calculate the significance of mediation (FDR-corrected p value cutoff ≤ 0.05). Mediation analysis indicated that CENPA completely mediated the effect of *LINC00261* on TMB, which may indicate a functional relationship between *LINC00261* and CENPA in coordinating DNA repair (Figure 4b). Although CENPA is not a canonical member of any DNA damage repair pathway (Supplementary Table 2) [41], there is previous work suggesting a role in the repair of double strand breaks, specifically via NHEJ [28,43]. In addition, GTF2H3, XPC and XPA, which are involved in the NER pathway [44], along with TOP2A, a crucial factor involved in proper DNA replication [45], partially mediated the relationship between *LINC00261* and TMB (Figure 4b and Supplementary Figure 3). Other DDR factors, including ATM and ATR, the NER factors GTF2H4, XPB, XPD, DDB1 and DDB2 and oncogenes such as KRAS and BRAF, showed a *LINC00261*-independent association to TMB (Figure 4b and Supplementary Figure 3).

In order to validate our bioinformatic findings that *LINC00261* alters DNA repair response, we treated A549 cells with or without stable sh*LINC00261* levels with platinum-based chemotherapeutic agents that work by increasing DNA damage resulting in genomic instability due to inactive repair mechanisms present in cancer cells. Both A549 shScrambled and A549 sh*LINC00261* cell lines underwent a dose response course with cisplatin, the parent platinum-based chemotherapeutic agent, as well as derivatives oxaliplatin and carboplatin, all of which have shown efficacy in treating LUAD and can be included in the standard of care for disseminated LUAD [46,47]. A549 cells with stable sh*LINC00261* knockdown were more sensitive to cisplatin (Figure 4c). The LD50 for A549-shScrambled was 5.80 μM, whereas the LD50 for A549-sh*LINC00261* was 0.06 μM. No significant differences in viability were observed for the derivatives carboplatin nor oxaliplatin (Figure 4d,e). Standard of care chemotherapy is often a combination of a platinum-based agent as well as the microtubule-disrupting paclitaxel [48]. To determine if *LINC00261* expression levels altered response to paclitaxel, we also treated A549 cells with stable sh*LINC00261* with paclitaxel and saw no significant difference in susceptibility (Figure 3f). The susceptibility towards cisplatin due to *LINC00261* silencing suggests that *LINC00261* exerts a protective effect from damage caused by cisplatin. To investigate whether *LINC00261* is involved in the response towards cisplatin damage, we treated cells with low dose (1 to 4 μM) of cisplatin for 3 days. Expression of *LINC00261* increased when A549 cells were treated with cisplatin at 2 and 4 μM, with a significance of 0.05 and 0.0051, respectively, compared to no treatment control (Figure 4e).

### 3.5. LINC00261 activates MHC Class II genes and alters immune cell composition in LUAD

The relationship observed between *LINC00261* and TMB suggested that neoantigen presentation may be altered in LUAD depending on *LINC00261* status. To determine whether there is an accumulation of neoantigens in LUAD patients with silenced *LINC00261* expression, we stratified TCGA-LUAD tumor samples based on *LINC00261* expression levels (cutoff levels were FPKM ≥ 1) and utilized NetMHCpan to determine neoantigen expression [49,50]. The neoantigen load for a given sample corresponds to the number of neoantigenic peptides due to single nucleotide variants (SNV) which is predicted to bind to the expressing HLA typing [50,51]. We observed that LUAD tumors with silenced *LINC00261* demonstrated lower numbers of predicted neoantigens (Figure 5a).

**Figure 5.**
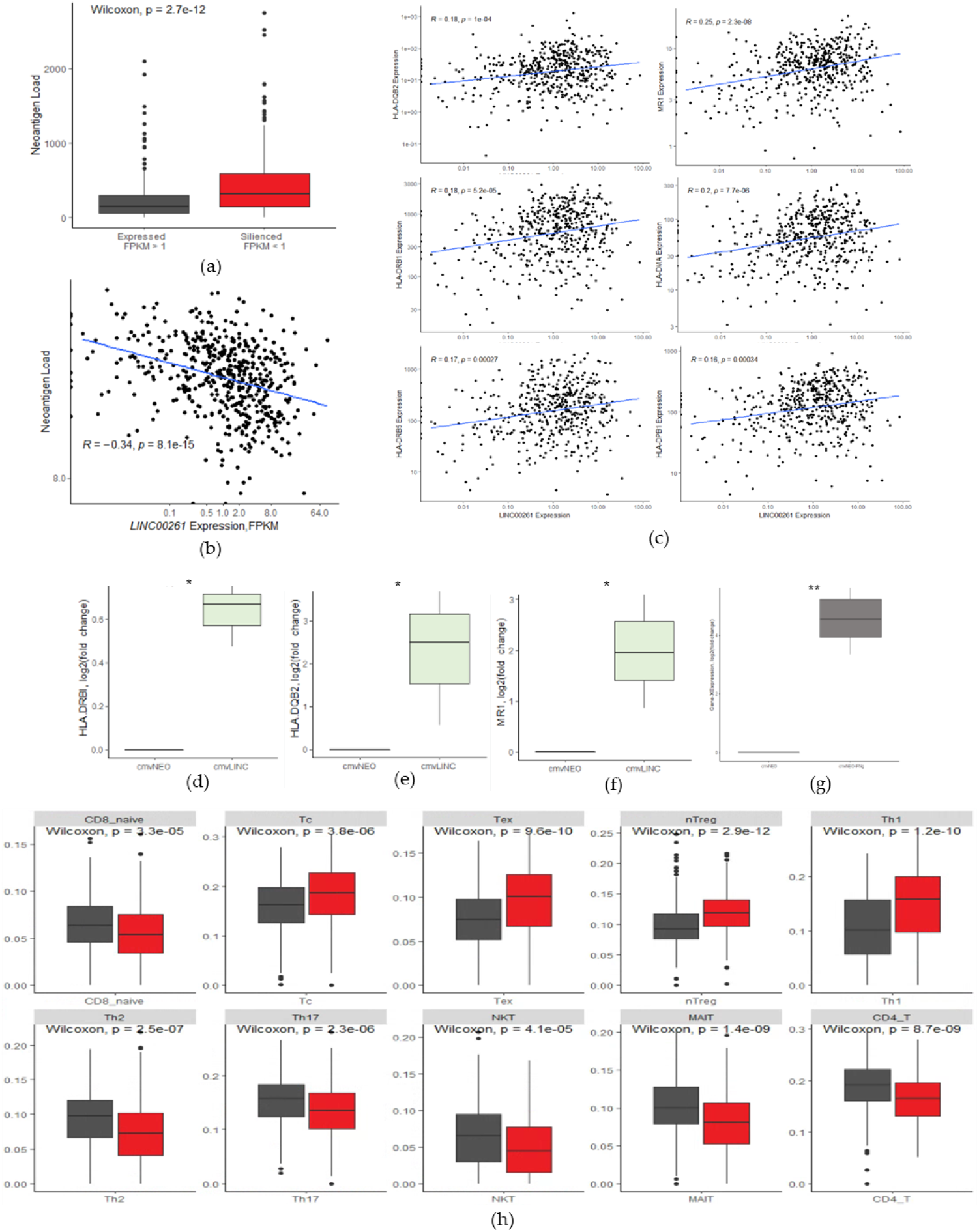
Expression of *LINC00261* confers with changes in MHC-class II gene expression composition of the tumor-immune microenvironment. (a) Amount of neoantigen load in TCGA-LUAD samples with expressing or silenced *LINC0261* expression. (b) Scatter plot of neoantigen load compared to *LINC00261* expression, in FPKM-UQ. (c) Scatter plot of MHC genes expression compared to *LINC00261* expression, in FPKM-UQ. MHC genes include, from counterclockwise, starting from the top right, *HLA-DQB2, HLA-DRB1, HLA-DRB5, HLA-DPB1, HLA-DMA, MR1*. (d-f) RTq-PCR of MHC genes between H522-CMV-NEO and H522-CMV-*LINC00261*. (g) RTq-PCR of *LINC00261* expression of H522 cell lines when treated with 50 ng/mL of INFγ or without (vehicle control of water). (h) Fraction of immune cells present in TCGA-LUAD samples, based on bulk RNA-Seq using xCell immune cell caller. Grey-Samples expressing *LINC00261* red-samples with silenced *LINC00261*.

**Figure 6.**
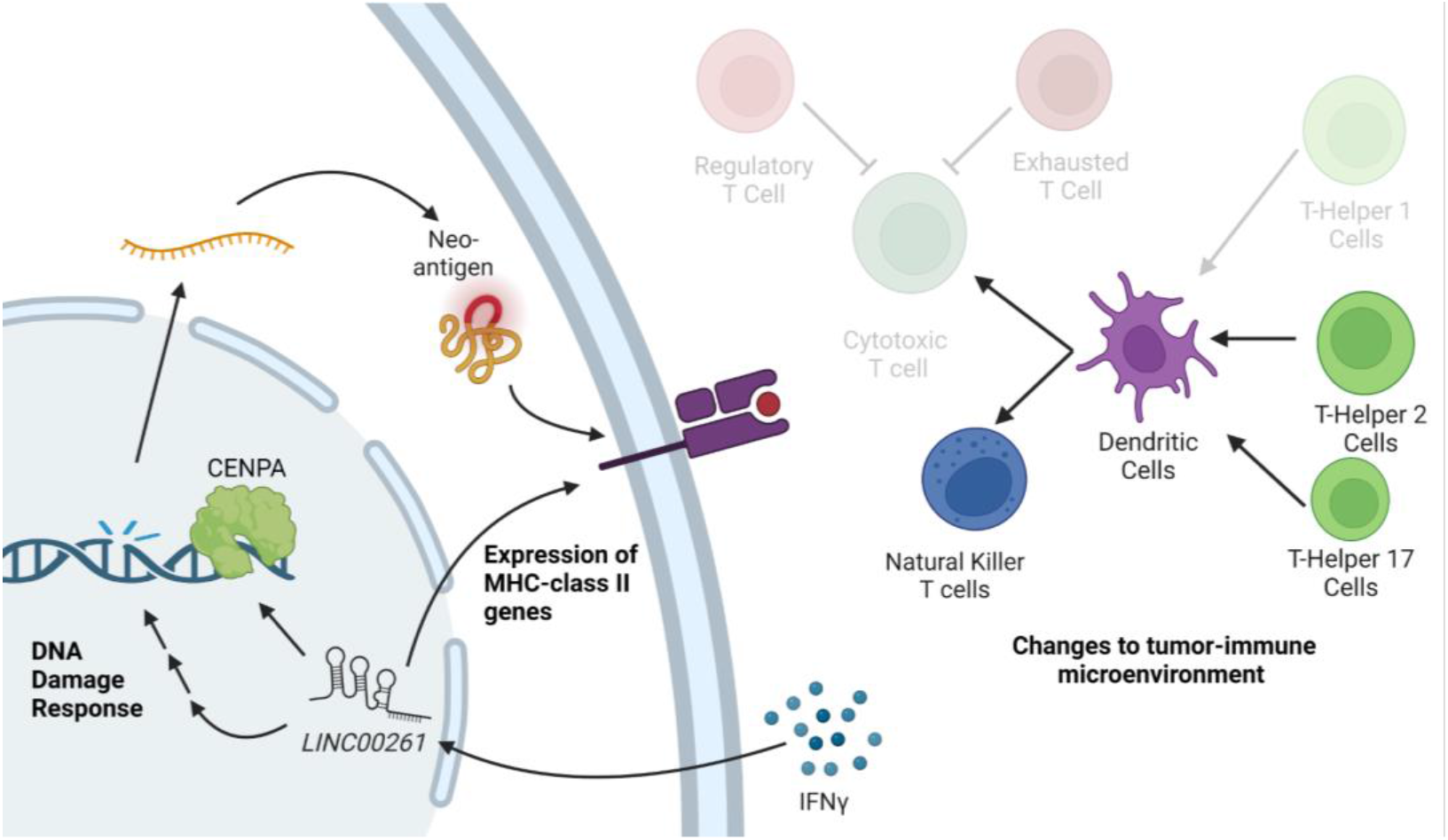
Mechanism of action by which *LINC00261* expression results in alterations to DNA damage repair, MHC-class II gene expression, and changes in tumor-immune microenvironment. Non-transparent immune cells are those which are in higher abundance in tumors expressing *LINC00261*. Transparent immune cells are those which are in lower abundance in tumors expressing *LINC00261*.

We next sought to understand the relationship between *LINC00261* and MHC Class II pathway genes. We examined correlations between *LINC00261* expression and pathways involved in antigen presentation, mainly the MHC Class I and Class II pathways, both of which are of importance in activating the anti-tumor immune response [52,53]. We identified multiple genes within the histocompatibility complex (MHC) Class II, including but not limited to, *HLA-DRB1, HLA-DRB5, HLA-DPB1* and *HLA-DMA* with a significant p-value of 5.2×10^-5^, 2.7×10^-4^, 3.4×10^-4^, 7.7×10^-7^, respectively (Figure 4B). A complete list of MHC genes correlating to *LINC00261* in LUAD is found in Supplementary Table 3. In addition, some MHC Class I genes were found to be correlated with *LINC00261* including *MR1* and *HLA-E* (Supplementary Table 3).

To validate this association, we investigated whether the re-introduction of *LINC00261* in LUAD cell lines altered the expression of genes encoding MHC components *in vitro*. The H522 LUAD cell line was stably transfected constitutively express *LINC00261* (H522-CMV-*LINC00261*) or empty vector control. In *H522-CMV-LINC00261* we observed an increase in expression of multiple MHC-Class II genes, including *HLA-DRB1* and *HLA-DQB2* (Figure 5c), consistent with the correlations between MHC Class II genes and *LINC00261* observed in the TCGA cohort.

In cancer cells, the expression of MHC-II plays a critical role in the recognition of a tumor to the immune system, resulting in better prognosis and improved response to immunotherapy [53,54]. The canonical mechanism by which MHC-II remains expressed is via IFNγ induction, where IFNγ-mediated activation of JAK1/JAK2 leads to expression of the transcriptional master regulator class II transactivator (CIITA) [53]. Because of our observed association between *LINC00261* and MHC-Class II gene expression, we investigated whether *LINC00261* is also among the downstream targets of IFNγ. We treated H522 cells with recombinant IFNγ at 1500 IU/mL for 24 hours, where we saw an activation of *LINC00261* expression, in which the gene is silenced in these LUAD cell lines (Figure 5d).

A crucial component of the anti-tumor immune response is the presence of pro-inflammatory and tumoricidal lymphocytes, such as natural killer cells and T-helper cells [55–57]. The absence of MHC Class II gene expression can influence the presence of immunological active cells within the tumor microenvironment [53]. To determine whether these changes in MHC Class II genes and predicted antigen presentation due to *LINC00261* expression corresponds with changes in the tumor-immune microenvironment, the fraction of immune cells present in TCGA-LUAD samples were calculated based on bulk RNA-Seq using xCell immune cell caller (Figure 5e) [58]. TCGA-LUAD patients expressing *LINC00261* had elevated levels of T-helper cells and natural killer T cells (Figure 5e), of which are key components of anti-tumor immune response [59]. In addition, an increase in exhausted T and regulatory T cells was observed in patients with silenced *LINC00261* cells which hinder the antitumor immune response [59].

## 4. Discussion

The accumulation of mutations is a key driving force in progressing cells into early stages of tumorigenesis [60]. Human cells have multiple mechanisms by which to protect genomic DNA, however, exposure to environmental mutagens, random mutations, and/or genetic inheritance can render critical DNA repair genes inactive, resulting in the accumulation of more mutations [5]. Although the field has primarily focused on protein coding genes whose loss of function results in diminished DNA repair capabilities, non-coding RNAs are emerging as genes of interest [17]. This is especially important as, in the case of LUAD, about 20-30% of patients lack the targetable mutated protein coding genes for targeted therapies, such as mutated KRAS and EGFR [61]. Results of this study suggest that *LINC00261* is a lncRNA whose expression is of clinical relevance, where LUAD cell lines with silenced *LINC00261* expression are more susceptible to cisplatin.

Beyond gene level biomarkers, immunological level biomarkers have been used to identify immunologically hot LUAD tumors that would benefit from immune checkpoint inhibitors [13]. While the use of proxy markers for neoantigen presentation, such as TMB, have aided in identifying patients who will respond to immunotherapy agents [6,62], there are still patients with high mutational burden that are not responsive to immune checkpoint inhibitors [63]. In addition, multiple studies have shown that mutational burden alone is not sufficient to serve as a biomarker for immunotherapy [64], pointing to the need of expanding the list of molecular biomarkers for more robust targeting of therapies.

The loss of *LINC00261* expression demonstrates several changes in pathways that are crucial indicators of successful treatment to therapy. Previous studies have shown that *LINC00261* is associated within the DNA damage signaling [18]. This study expands the understanding of how *LINC00261* silencing in LUAD is involved in an increased amount of TMB, changes in DNA repair pathways, and cisplatin-mediated susceptibility to the DNA damaging agent.

In addition, it is likely that the change in mutational accumulation influences the tumor microenvironment. *LINC00261*-silenced tumors demonstrated a more unfavorable anti-tumor immune response, with a higher number of exhausted and regulatory T-cells, along with a decrease in cytotoxic and helper T cells (Figure 5h). Taken together, we have identified a lncRNA, *LINC00261* whose expression status coincides with clinically relevant changes to the tumor. Our study points towards clinical relevance in identifying expression status. In identifying clinically relevant lncRNA in LUAD, our work demonstrates potential applications of non-coding RNAs as molecular biomarkers in expanding targeted therapeutics.

## Author Contributions

Conceptualization - JC, AE, TM, CNM; methodology - JC, CNM; formal analysis - JC, TX, DJM; resources - CNM; data curation - JC, TX, SJ, MN, SB, SJ; writing original draft preparation - JC; writing, review and editing - JC, CNM.; visualization - JC, TX; supervision - CNM; funding acquisition - CNM. All authors have read and agreed to the published version of the manuscript.

## Funding

This research was funded by a Research Scholar Grant to C. Marconett [RMC-RSG-20-135-01], an IDEA award from the Department of Defense Lung Cancer Program [W81XWH-21-1-0231], the Baxter Foundation, and the Departments of Translational Genomics and Surgery at the Keck School of Medicine. J. Castillo was supported by a Dean’s Scholar award from the Keck School of Medicine. S. Joseph was supported by the CaRE2 U54 postbaccalaureate premedical training program [CA233396]. This project was supported administratively and technically by the Norris Comprehensive Cancer Center core grant [P30 CA014089].

## Institutional Review Board Statement

Analysis of publicly-available human data was approved under exempt IRB #HS-21-00160. Utilization of human cell lines for RNA analysis and chemotherapeutic response was approved under exempt IRB #HS-20-00577.

## Informed Consent Statement

Not applicable

## Data Availability Statement

TCGA data utilized in this study is publicly available through the Genomics Data Commons (GDC) portal (https://portal.gdc.cancer.gov).

## Acknowledgments

We would like to thank Ite Offringa, PhD for discussion of topics relating to the manuscript, Matt Michael, PhD for discussion of DDR signaling mechanisms and crosstalk, and John Carpten, PhD for discussion of TMB relevance for immunotherapeutic responses.

## Conflicts of Interest

The authors declare no conflict of interest. The funders had no role in the design of the study; in the collection, analyses, or interpretation of data; in the writing of the manuscript; or in the decision to publish the results.

## Supplementary Figures

**Supplementary Figure 1.**
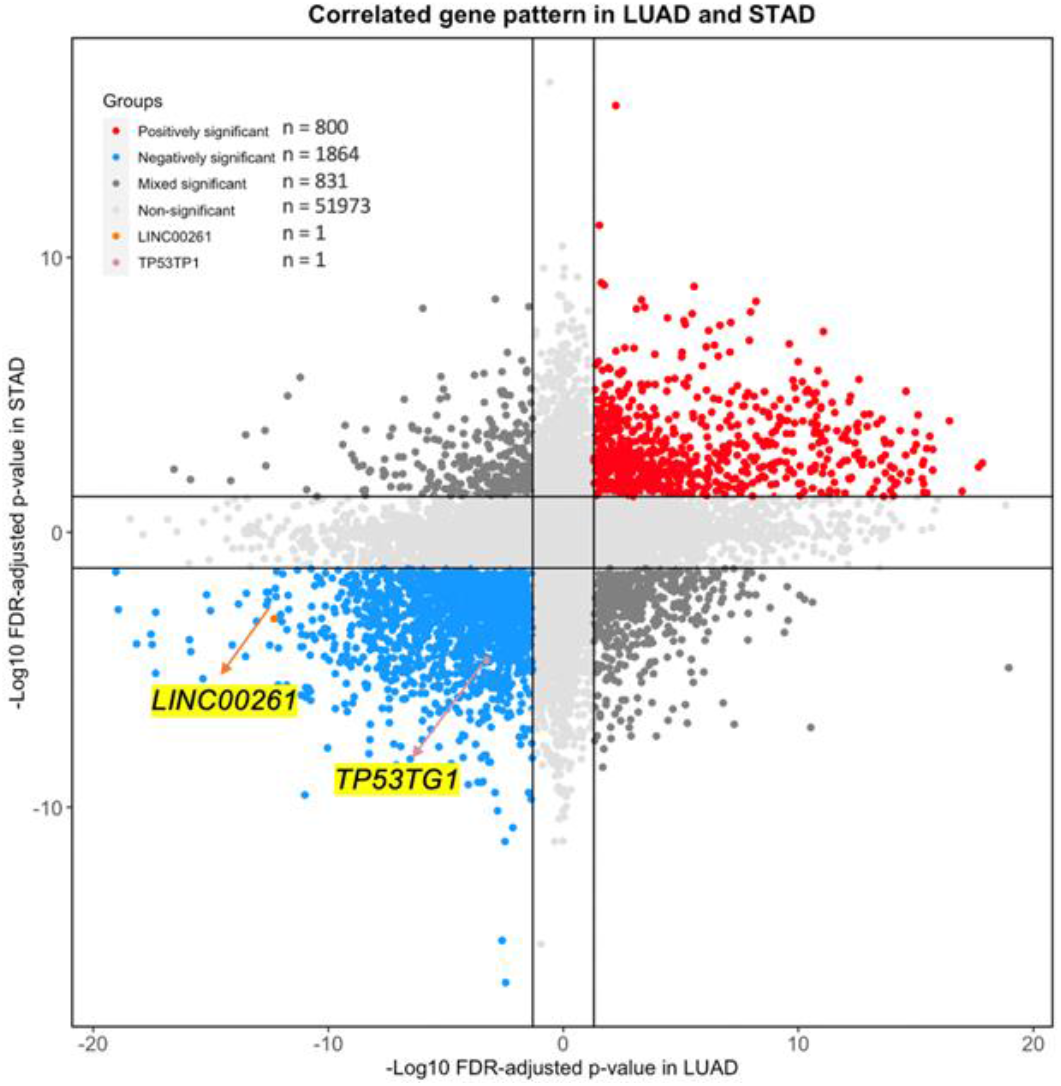
Red points = dual-positively TMB-correlated genes, blue points = dual-negatively TMB-correlated genes, dark gray points = TMB-correlated genes with opposite correlation values in LUAD and STAD, light gray points = genes which were not significantly correlated to TMB in LUAD and STAD. *LINC00261* was marked as the orange point, *TP53TG1* was marked as the pink point.

**Supplementary Figure 2.**
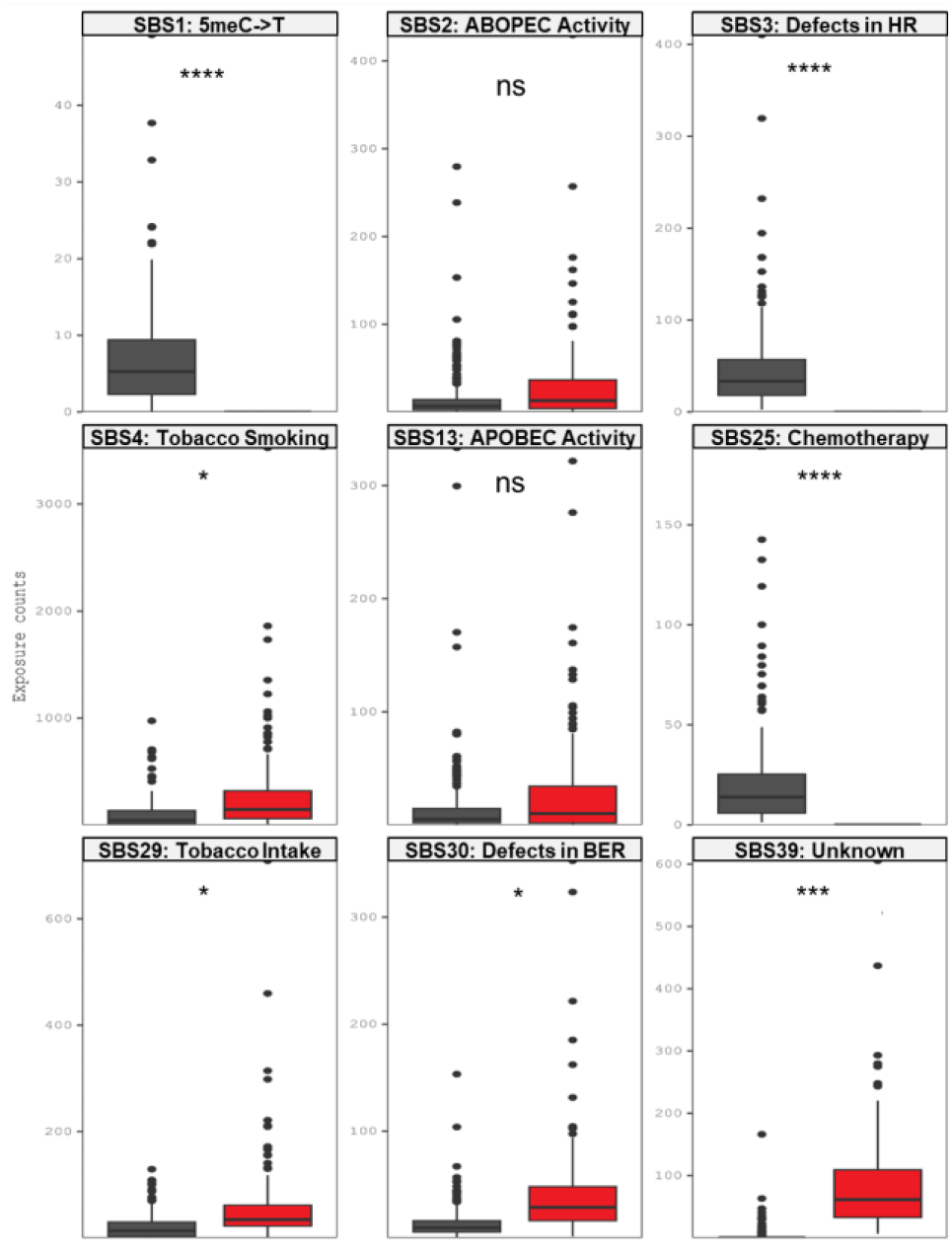
Distribution of exposure counts for the nine SBS mutational signatures between the *LINC00261*-expressing and *LINC00261*-silenced groups.

**Supplementary Figure 3.**
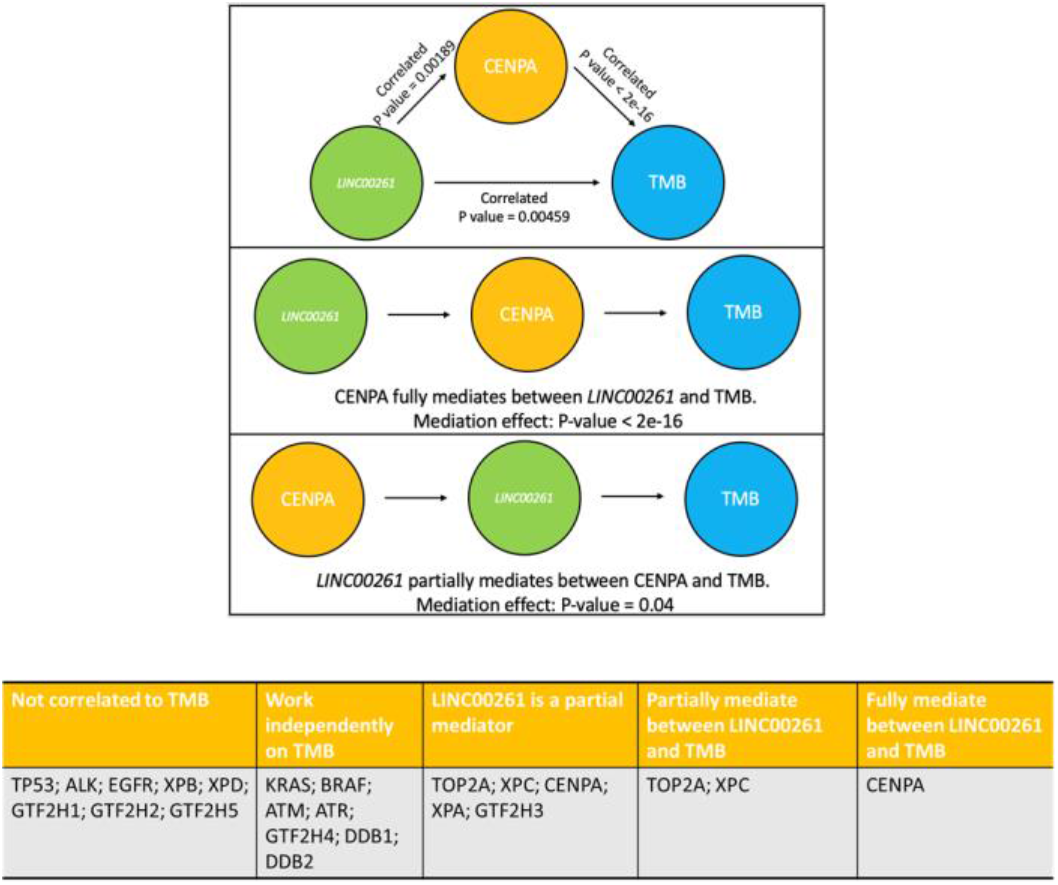
Mediation analysis results between *LINC00261* TMB, and tested mediator factors.

## Notes

### Competing Interest Statement

The authors have declared no competing interest.

